# Quantitative blood flow estimation *in vivo* by optical speckle image velocimetry

**DOI:** 10.1101/2021.03.25.437094

**Authors:** Muhammad Mohsin Qureshi, Yan Liu, Khuong Duy Mac, Minsung Kim, Abdul Mohaimen Safi, Euiheon Chung

## Abstract

Speckle based methods are popular non-invasive, label-free full-field optical techniques for imaging blood flow maps at single vessel resolution with a high temporal resolution. However, conventional speckle approach cannot provide an absolute velocity map with magnitude and direction. Here, we report a novel optical speckle image velocimetry (OSIV) technique for measuring the quantitative blood flow vector map by utilizing particle image velocimetry with speckle cross-correlations. We demonstrate that our OSIV instrument has a linearity range up to 7 mm/s, higher than conventional optical methods. Our method can measure the absolute flow vector map at up to 190 Hz without sacrificing the image size, and it eliminates the need for a high-speed camera/detector. We applied OSIV to image the blood flow in a mouse brain, and as a proof of concept, imaged the real-time dynamic changes in the cortical blood flow field during the stroke process *in vivo*. Our wide-field quantitative flow measurement OSIV method without the need of tracers provides a valuable tool for studying the healthy and diseased brain.

## 1. Introduction

Although the human brain occupies only 2% of body mass, it consumes 20% of the oxygen and nutrients supplies (e.g., glucose) of the body [1]. The local cerebral blood flow (CBF) regulation via brain vasculature plays a crucial role in transporting oxygen and glucose to the location of the neural activity [2–4]. In clinical and research settings, measuring blood flow in cerebral vessels is of paramount importance for understanding cerebral metabolism and cerebrovascular pathophysiology [5–7].

Various tools have been developed to measure and monitor the spatiotemporal dynamics of CBF, as CBF regulation provides the key to unravel the coupling of local blood flow control in response to the nearby neuronal activity [8–10]. Optical methods for CBF measurements fall under three categories: 1) Doppler based methods, such as laser Doppler flowmetry, Doppler optical coherence tomography, and photoacoustic Doppler velocimetry [11–15]. These methods are quantitative, but wide field-of-view measurement at a high frame rate is limited. 2) Red blood cell (RBC) tracking measurements, such as intravital multiphoton laser scanning microscopy (MPLSM), confocal laser scanning microscopy (CLSM) and holographic phase microscopy [16–19]. However, MPLSM and CLSM measure a limited number of vessels because of their point scanning nature. The holographic phase approach is useful for thin samples. 3) Speckle-based methods, such as laser speckle contrast imaging (LSCI) [6,20] and multi-exposure speckle imaging (MESI) [21].

Speckle-based approaches require simple instrumentation and can provide a high spatial or temporal resolution for label-free, wide-field imaging of CBF that can be easily applied in clinics [22]. However, conventional LSCI methods provide only relative blood flow information which is unsuitable for longitudinal studies [6]. To resolve this fundamental issue, speckle imaging for quantitative blood flow measurement has become an active research area. Thus far, three studies laid the groundwork, which are frequency-domain laser speckle imaging (FDLSI) [23], laser speckle flowmetry [24], and multi-exposure speckle imaging (MESI) [21]. FDLSI involves a mathematical model incorporating Gaussian distribution and the Brownian distribution of particles for highly ordered motion and may not be accurate within *in vivo* samples. Laser speckle flowmetry introduced a scaling factor for the correction of the multiple scattering in the microcirculatory vessels; however, the scaling factor has to be recalculated for different samples. MESI uses different camera exposure times with a time-gating approach of laser illumination for obtaining an enhanced quantitative flow measurement. Recently, the dynamic light scattering imaging model [25] showed improvement in LSCI and MESI models and demonstrated the cortex field-of-view of a whole mouse brain with an accurate estimation of speckle correlation time via their speckle intensity temporal autocorrelation function. However, all these methods require mathematical modeling or calibration for quantitative flow measurement. Conventional laser speckle imaging methods have an inherent limitation to compare blood flow changes even within the same animal/human blood vessels or with control blood vessels. Simply the outcome is not quantitative and thus incomparable. In contrast, our approach allows quantitative comparison of the same subject’s blood flows over time and enables the comparison of treatment efficacy for vascular diseases in a longitudinal fashion, a key capability for preclinical and clinical studies.

On a different front, particle image velocimetry (PIV) is a technique that utilizes sequential images of moving particles to extract average speed and direction [26,27]. However, the tracer seeding problem limits the use of the conventional PIV technique for *in vivo* applications. RBCs have been reported to be used as tracer particles of PIV with wide-field microscopy. However, this approach was limited to highly transparent vessels such as mesentery vessels for *in vivo* studies [28] or vessels found in the transparent chicken embryo [29]. The PIV technique was recently used with confocal microscopy to estimate the blood flow quantitatively in the sub-surface regime [30]. However, this study still achieved a limited velocity range of up to 1 mm/s at 180 fps and 200 μm/s at 30 fps, because of a point-by-point scanning method.

In this paper, we present a simple technique by combining the PIV method with laser speckle image analysis for wide-field, quantitative RBC velocity measurement. The technique is termed optical speckle image velocimetry (OSIV). OSIV is analogous to fluorescent speckle microscopy (FSM) [31]. However, for FSM, speckles were formed by the random association of fluorophores with a macromolecular structure. In contrast, in OSIV, speckles were developed by the interaction of the particle with a coherent laser light. As in the PIV approach, temporal cross-correlation of consecutive speckle images provides the OSIV velocity fields. We validated the approach proposed herein by using, moving glass diffuser, flowing RBCs in a microfluidic channel *in vitro* and by measuring the cortical blood flow of a mouse *in vivo*. Furthermore, we demonstrated the application of OSIV with a preclinical ischemic stroke model via photothrombosis (PT). To the best of our knowledge, the technique proposed herein is the first to use laser speckles to provide a quantitative dynamic velocity map of an *in vivo* mouse brain and a cortical stroke model that allows longitudinal assessment of potential therapeutic interventions objectively.

## 2. Methods

### 2.1. Working principle of OSIV

When light from a coherent source interacts with a material, a granular pattern with high contrast known as laser speckle is obtained because of multiple scattering of light by the particles in the medium [32]. A speckle pattern remains unchanged over time if the laser light interacts with a static medium. However, if the sample is not static (e.g., blood flowing in a living tissue), the laser speckle pattern would be dynamic, as shown in our previous studies [33–36].

The OSIV technique is inspired by PIV, which requires a minimum of two consecutive images [27]. OSIV is based on the cross-correlation of speckle images and the OSIV algorithm is illustrated in Fig. 1. Suppose that two successive images were captured with a camera, and the images are divided into small probing windows *I*_1_(*x, y*) and *I*_2_(*x, y*). We compute the correlation of *I*_1_(*x, y*) and *I*_2_(*x, y*) by using the cross-correlation theorem. Specifically, we take the Fourier transform of the probing windows and multiply the Fourier transform of the first window *F*_1_(*u, v*) with the conjugate of the second 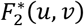. After taking the inverse Fourier transform, we obtain the correlation map *c_map_* for a single realization.

**Fig. 1.**
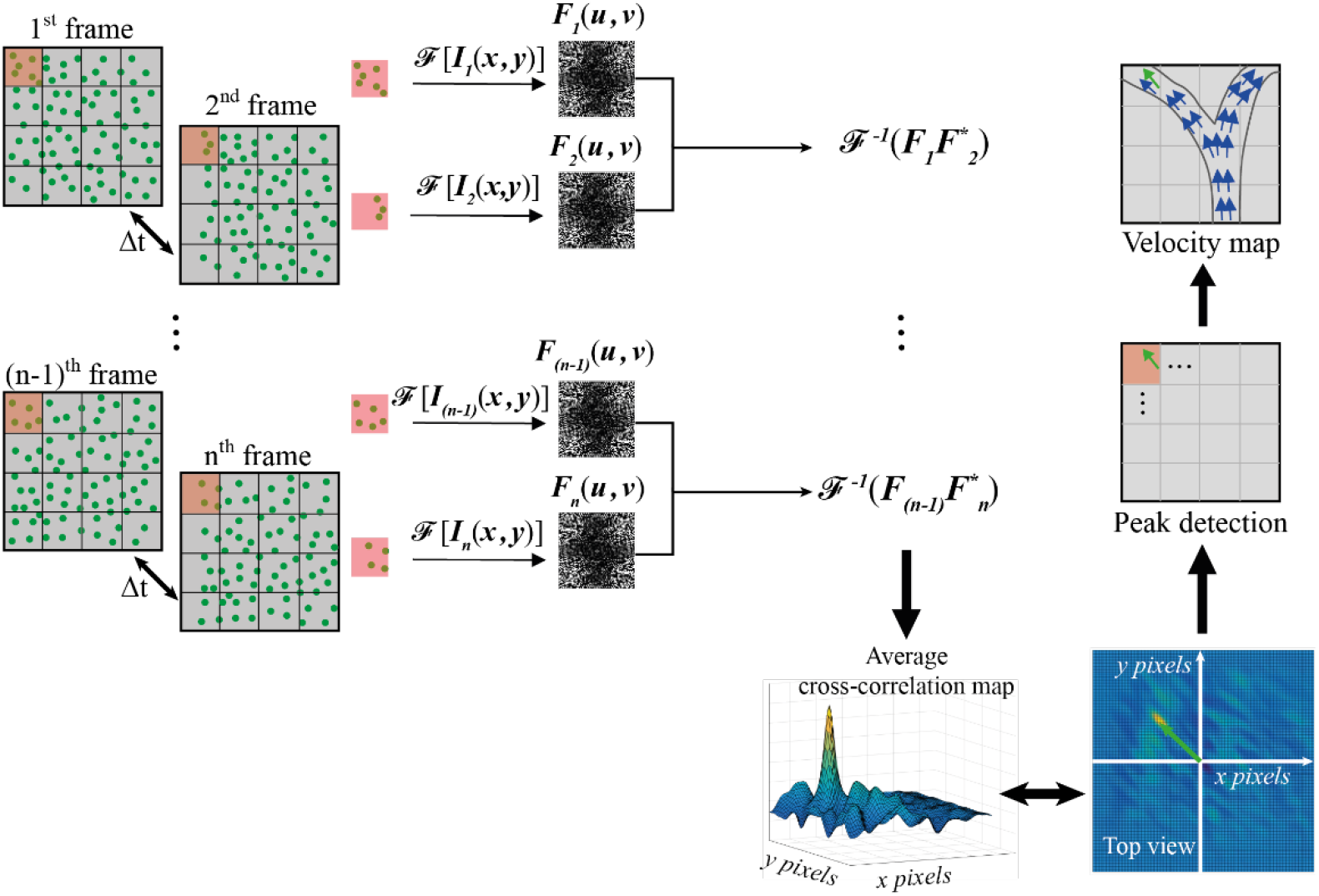
Working principle of optical speckle image velocimetry (OSIV). Consecutive frames are taken at a time interval of ‘*Δt*’ On each image pair (i.e., the (n−1)-th frame and the n-th frame), a window (red square) is selected at the same location and Fourier transform is performed on the windowed region 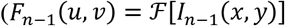 and 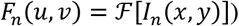. We multiply *F*_*n*−1_(*u, v*) with the conjugate of *F*_*n*_(*u, v*), and then take the inverse Fourier transform of the product to obtain the cross-correlation map. These steps are repeated for ‘n’ frames; thereafter, we take the average of the cross-correlation maps and find the peak to get the displacement d, from which we obtain the velocity v at this window location by v = d/*Δt*. We obtain the velocity map of the field of view by shifting the window location and repeating the aforementioned procedure at each window location.

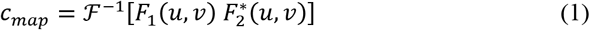

We averaged the correlation maps over *n* realizations. In this study, *n* = 4 was used to reduce the noise and obtain the OSIV velocity map. From the average correlation map, the peak location can be measured, which gives the displacement of the particles during the time interval. We estimated the flow speed at the probing window location by dividing the displacement with the reciprocal of the camera frame rate. Then, to obtain a velocity map of the field of view, we shifted the probing window pixel-by-pixel and repeated the aforementioned procedure at each window location, until the whole speckle image was scanned by the probing window completely.

A critical parameter to consider was the size of the probing window in the OSIV algorithm. Two factors must be considered while selecting the size of the probing window. First, the window size should be smaller or comparable to the region of interest (ROI). Second, the window size also defined the OSIV capability to measure the maximum velocity magnitude, that is, the bigger the window size, the higher is the speed that can be measured. If the ROI is defined in terms of *N × N* pixels, and the optical system has the magnification *OZF* (optical zoom factor), the linear length *l* of the probing window of *“m × m”* pixels in microns is calculated as follows:

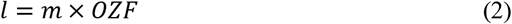

If the time interval between the two consecutive speckle images is Δ*t*, the maximum velocity that can be estimated by OSIV would be

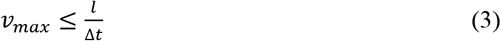

This inequality always holds, because in the OSIV technique, we consider a particle with a size ranging from 3 to 5 pixels as a speckle. Then, the spatial displacement between the consecutive probing windows should be less than the length *l*. In this study, a camera with a maximum frame rate of 190 fps was used with a probing window size of 64 × 64 pixels. For the moving diffuser and fluidic phantom with a 10× objective lens, the *OZF* value was 0.648 μm/pixel. However, for the *in vivo* experiment with a 20× objective lens, the *OZF* was 0.321 μm/pixel. Therefore, with the current optical parameters and probing window size, the maximum speed measured from OSIV was approximately 7.5 mm/s for the 10× objective lens and 3.7 mm/s for the 20× objective lens.

### 2.2. Optical implementation

The detailed optical setup for OSIV is shown in Fig. 2. Speckles were formed by a green laser of wavelength 532 nm (PSU-III-LCD, Changchun New Industries Optoelectronics Technology Co., Ltd., China). To maintain a wide field-of-view and prevent speckles from smearing, we used an acousto optical modulator (AOM) (AOM-505AF1, IntraAction Corp. Bellwood, USA) for optical chopping of light. The laser beam was steered to AOM using a pair of mirrors, and only the first-order diffraction beam was passed through an iris. A function generator (4052 BK Precision Corp. California, USA) was connected to an AOM driver (ME-504, IntraAction Corp. Bellwood, USA) to control the AOM. The function generator produced square waves with a 20% duty cycle at the frequency of 190 Hz which is the maximum frame rate of the camera used in this study. With this AOM modulation, we achieved optical chopping of the continuous laser light. During the camera exposure time of *t_e_*, the modulated light by the AOM was on for the duration of *t_A0M_*. Therefore, the effective sample illumination time *t_s_* on the camera was the same as *t_A0M_* as shown in Fig. 2(a).

**Fig. 2.**
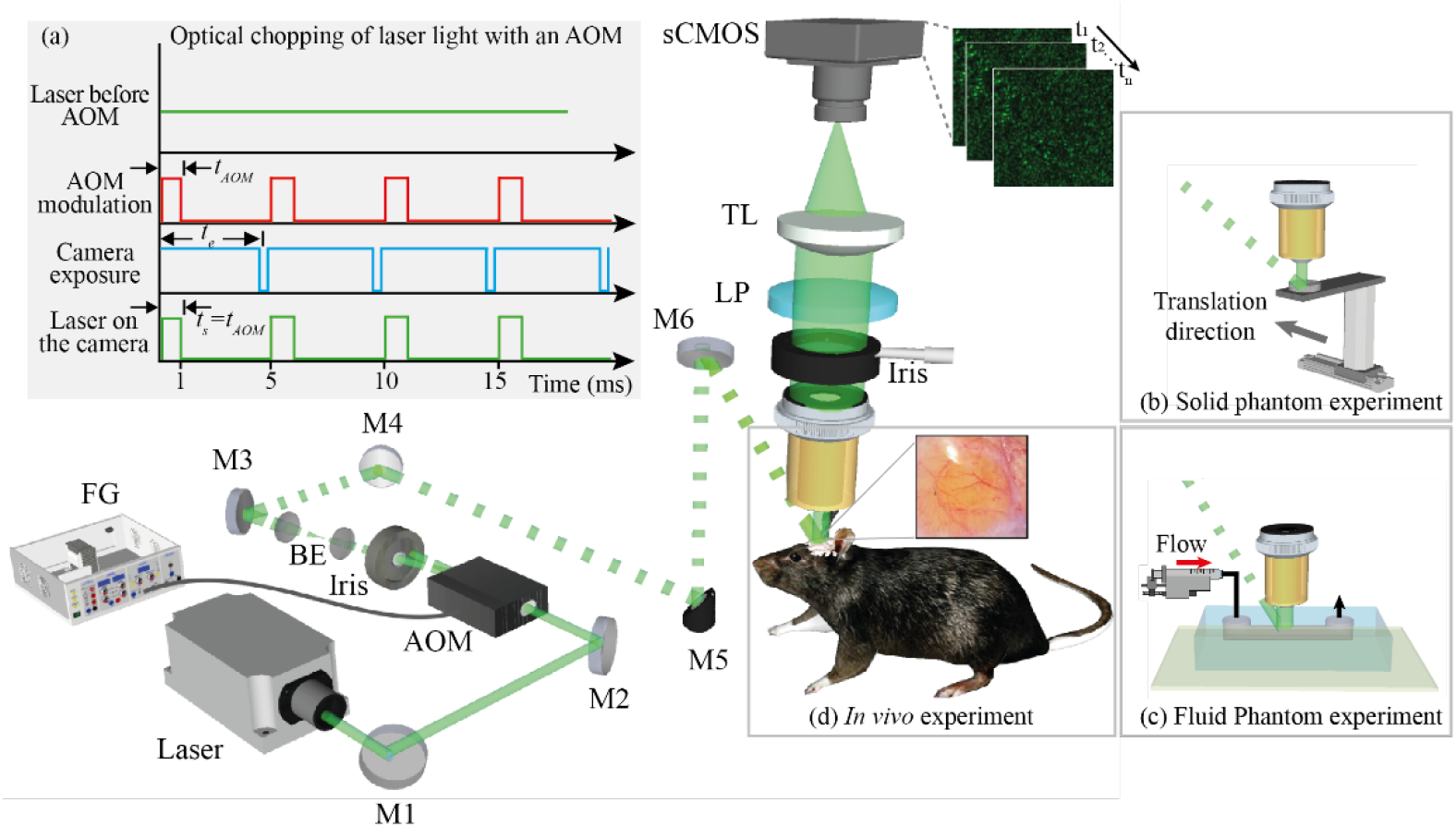
Experimental setup for OSIV. (a) Schematic of the setup. A 532 nm wavelength laser is used as a coherent source to obtain speckle images. Laser beam passes through an acousto optical modulator (AOM) which is connected with a function generator for optical chopping of light at the same rate with the camera frame rate. The reflected light from the sample is collected with an objective lens and it passes through an iris, polarizer, tube lens and is finally captured by a camera which records a stack of speckle images. BE: beam expander, FG: function generator, M1 to M6: mirror, LP: linear polarizer, TL: Tube lens. Inset of (a) illustrates optical chopping of the laser light with the help of the AOM and function generator. ‘*t_AOM_*’ is the AOM modulation time, ‘*t_e_*’ is the exposure time of the camera and ‘*t_s_*’ is the time interval in which sample is shinned with the laser light. (b) Setup for the *in vivo* experiment. (c) Setup for the solid phantom experiment. A diffuser is translated by a motorized stage. (d) Setup for the fluid phantom experiment. Red blood cells steadily flow through the microfluidic channel with the help of a syringe pump.

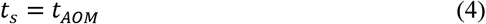

The value of *t_A0M_* was equal to 1.05 ms, and the minimum exposure time *t_e_* of the camera was set to 5.26 ms with an ROI size of 512×512 pixels. The AOM-modulated beam was expanded 1.5 times by a pair of convex lenses. The average power of the modulated laser beam on the sample was 3.5 mW. For the *in vivo* case, speckle imaging and white light imaging were performed sequentially.

The detection part consisted of an objective lens, an iris, a linear polarizer, a tube lens, and an sCMOS camera An objective lens of 10× magnification (RMS10X, NA 0.25, Olympus) was used for moving the glass diffuser and *in vitro* sample imaging. However, for the *in vivo* experiment, an objective lens of 20× magnification (PLN20X, Olympus) and NA of 0.40 was used. The reflected light from the sample was passed through an iris that increased the speckle size, a linear polarizer for improving the speckle contrast, and a tube lens with a focal length of 180 mm. Finally, the light was captured by an sCMOS camera (Neo 5.5 sCMOS, Andor Technology Ltd., Belfast, UK), which ran at its maximum frame rate of 190 fps. The iris was adjusted so that the speckle size on the camera was ~3 × 3 pixels to satisfy the Nyquist criteria for sampling the speckle. For all the speckle images, the speckle contrast was always greater than 0.5.

## 3. Results

### 3.1. Proof-of-principle with a moving diffuser

To validate the OSIV results, we designed a proof-of-principle experiment. With a moving glass diffuser illuminated by coherent laser light, we acquired a series of speckle images, as shown in Fig. 3(a). The raw speckle images were used to generate the OSIV maps for the movement speed of 2, 3, 4, 5, 6, 7, and 8 mm/s. A single representative OSIV map for each speed of 2, 4, 7, and 8 mm/s is shown in Figs. 3(b)–(e), and these maps are overlaid on one speckle image of the corresponding speeds. The length of the white arrow represents the speed, and the arrowhead shows the direction of movement of the diffuser.

**Fig. 3.**
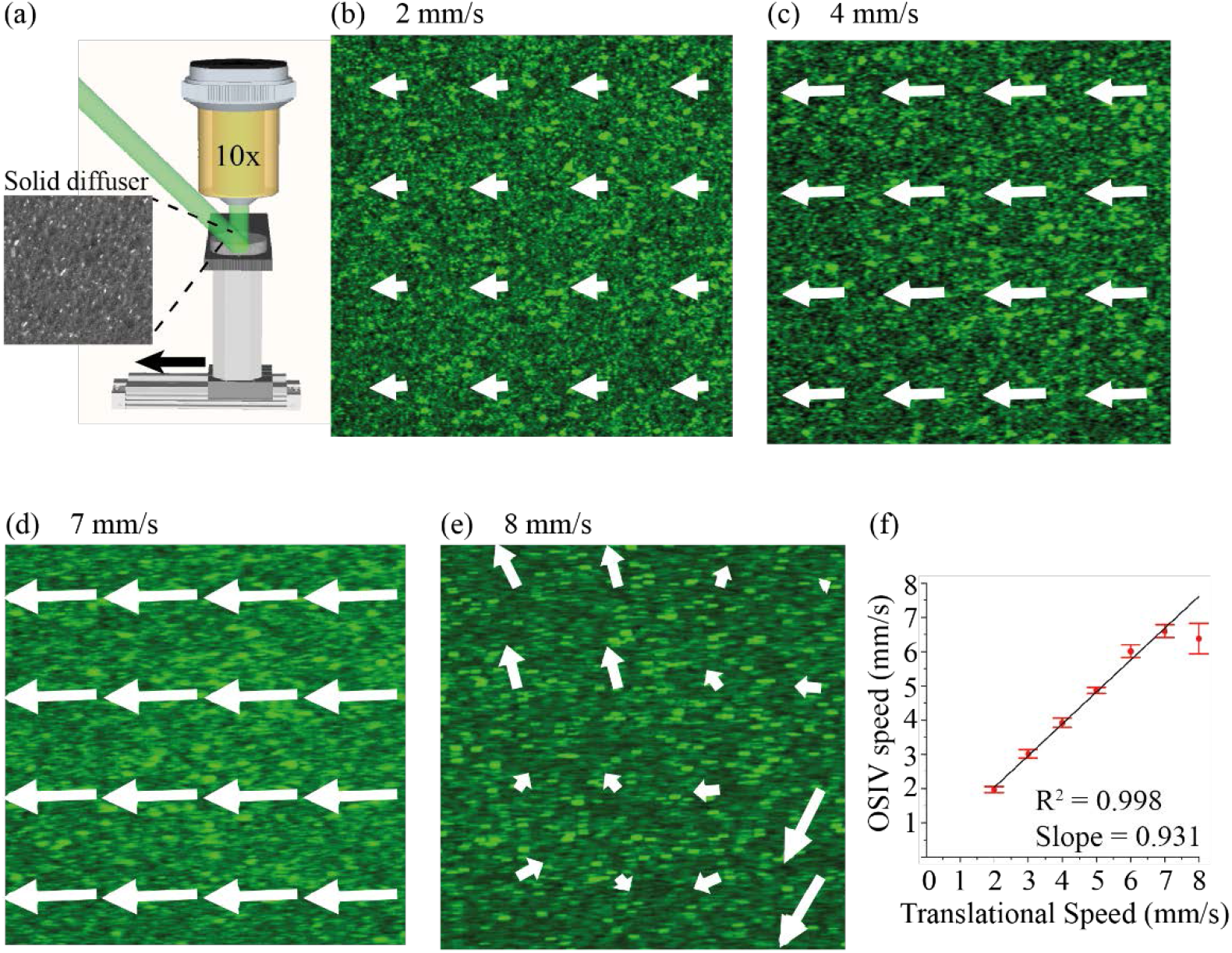
OSIV of a moving diffuser. (a) Schematic of a solid diffuser moved by a motorized stage from right to left and a 10X magnification objective lens for collecting the reflected light. (b)–(e) OSIV measured velocity map when the diffuser was moved with a constant speed of 2, 4, 7, and 8 mm/s, respectively. Image size: 330 μm across. (f) Relationship between the OSIV measured speed and the motorized stage speed. For each speed the number of trials was 10 (i.e., n = 10). The red data points show the mean value and standard deviation (SD) while the black line is a linear fit with R^2^ of 0.998 and a slope of 0.931. The velocity direction obtained from OSIV matches with that of the motorized stage.

We compared the speed measured by OSIV with the preset speed of the diffuser in Fig. 3(f). The estimated speed from OSIV matched the preset speed up to 7 mm/s (slope = 0.931). The OSIV underestimates beyond this limit, which was predicted from the theory in equation (3). To exemplify the dynamics of OSIV, movies of OSIV maps for preset speed of 4 and 8 mm/s are presented in Supplementary Movie S1. At 4 mm/s, the velocity arrows showed the same directions and magnitudes as expected, while the OSIV map showed random magnitudes and directions at 8 mm/s.

### 3.2. In vitro demonstration with RBCs flowing in a microfluidic channel

After OSIV demonstration with a solid diffuser, we tested whether OSIV is applicable in a more realistic configuration with moving RBCs in a microfluidic channel mimicking a blood vessel. The illumination and detection setup was similar to the previous experiment, and the microfluidic sample is shown in Fig. 4(a). For each fluid (RBCs) speed of 0.74, 1.48, 2.22, 2.96, 3.7 mm/s, one OSIV map is shown in Figs. 4 (b)–(f). The direction of the arrowhead in the OSIV map matches with the RBC flow direction, and the length of the arrow shows the speed. The relationship between the speed measured by OSIV and the preset RBC speed (calculated from the syringe pump flow rate) is shown in Fig. 4(g). The OSIV results were in good arrangement with preset speed up to 3.7 mm/s, which agreed with the expectation from the theory (i.e., Eq. 3).

**Fig. 4.**
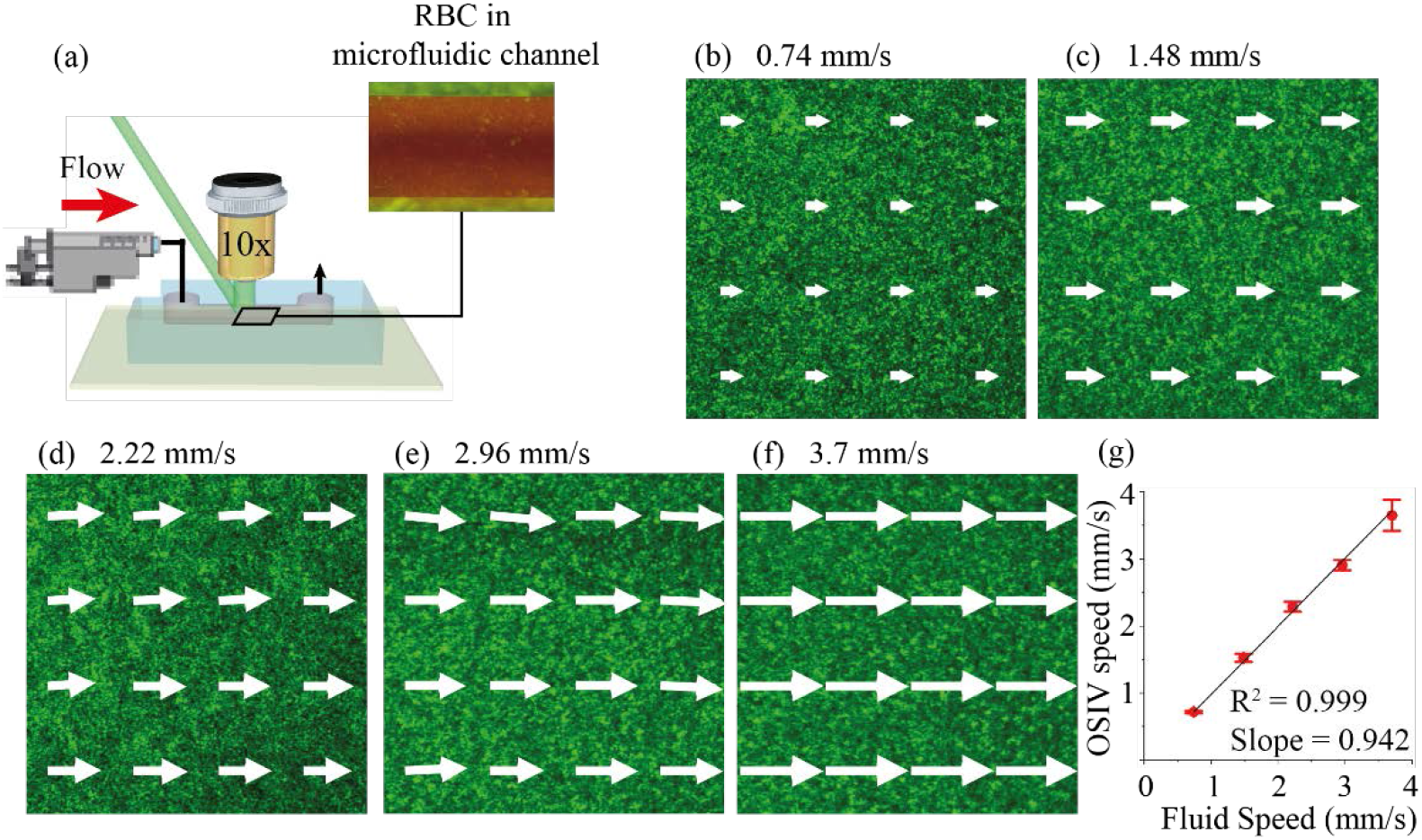
OSIV of flowing RBCs in a microfluidic channel. (a) Schematic of a fluidic phantom composed of washed-RBCs flowing at a steady rate from left to right in a microfluidic channel with the help of syringe pump, and an objective lens of 10X magnification for collecting the reflected light. (b-f) OSIV velocity maps when the speed calculated from the syringe pump was 0.74, 1.48, 2.22, 2.296 and 3.7 mm/s, respectively. Image size: 330 μm across. (g) Relationship between the OSIV measured speed and the preset speed of the fluid. For each speed the number of trials was 10 (i.e., n = 10). The red data points show the mean value and standard deviation while the black line is the linear fitting with R^2^ value of 0.999 and slope of 0.942.

### 3.3. In vivo demonstration of mouse cortical blood velocity map

After the solid diffuser and *in vitro* demonstration, we applied OSIV to generate the velocity map of a cortical blood vessel of a mouse *in vivo*. The selected ROI on the cranial window is shown in Fig. 5(a). The time-averaged OSIV maps of velocity field with direction and magnitude are shown in Figs. 5 (b),(c). These maps were produced by the average of 25 OSIV maps. The velocities in the blood vessel, obtained via OSIV, are directed downwards, as shown in Fig. 5(b). The enlarged inset shows the velocity vector field with the arrow length commensurate with the corresponding magnitude. A detailed map for the average speed is shown in Fig. 5(c). The flow profiles at two locations AA’ (red) and BB’ (blue), along with the calculated flow rates from OSIV maps, are shown in Fig. 5(d). The flow profiles were fitted with two parabolic curves. While the maximum velocities at the locations are different (0.35 and 0.42 mm/s), the calculated flow rates are similar without significant difference, following the principle of conservation of mass.

**Fig. 5.**
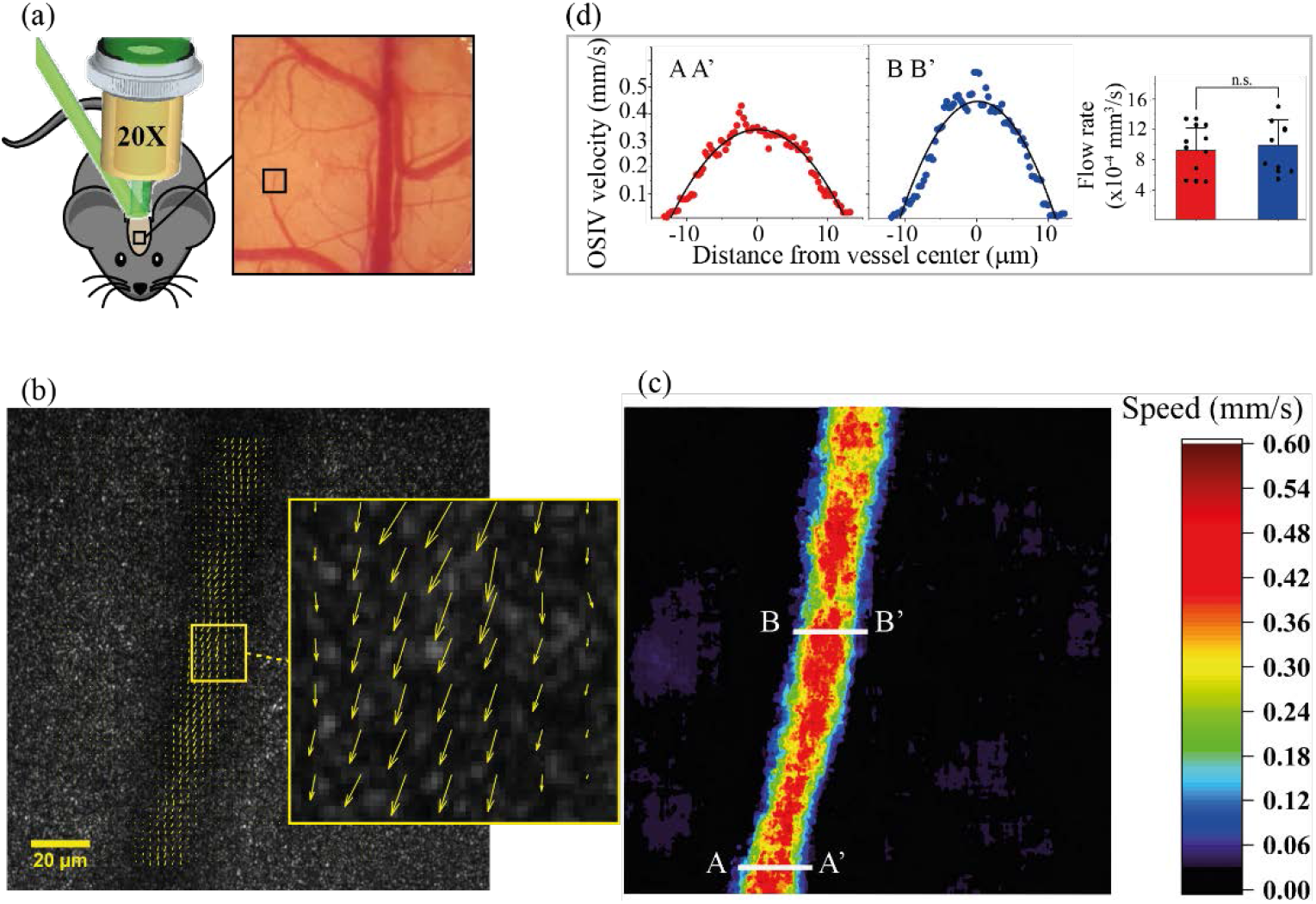
OSIV of blood flow in a mouse brain *in vivo*. (a) Schematic of the *in vivo* experiment and the location of imaging. (b) OSIV measured velocity map overlaid on the speckle image. We observe that blood flowed in the downward direction. (c) OSIV measured flow speed map. (d) the flow speed profiles along the two lines denoted by AA’ and BB’. The experimental data points were well fitted by a parabolic curve. The bar graph shows the flow rate measured at the two locations of AA’ and BB’ for 10 OSIV maps. We observe no significant difference between the flow rates at two locations. This exhibits the conservation of flow rate.

To confirm if the OSIV velocity values agree with another technique, we acquired two consecutive white light images for tracing individual RBCs tracked by the ImageJ plugin TrackMate. The histogram of RBC-tracing results falls within the range of 0.1 to 2.0 mm/s of the OSIV histogram (Supplementary Fig. S1). However, these two results were obtained sequentially.

### 3.4. Dynamic velocity field changes with a photothrombosis stroke model

Following the demonstration of *in vivo* cortical velocity mapping, we applied OSIV to a PT stroke model with additional optical fiber-based irradiation, as shown in Fig. 6(a). The photothrombotic light illumination was applied globally over the cranial window. A series of speckle images were acquired at −1, 1, and 4 min from the start of PT laser illumination for the pre-stroke, during stroke, and post-stroke conditions. We generated 25 OSIV maps for each condition. At the designated location ‘**P**’, the corresponding 25 OSIV velocity vectors are shown over Δ*t* time interval for the three conditions in Fig. 6(b). The velocities change dynamically in the direction and amplitude, reducing the average speed from *t_R_ = −1* to *t_R_ = 4* min. With the average OSIV maps, the RBC velocity vectors and magnitudes for the pre-stroke, during-stroke, and post-stroke condition are shown in Figs. 6(c),(f), Figs. 6(d),(g), and Figs. 6(e),(h), respectively. From the inset of the average OSIV map for each condition, it was observed that RBC flow speed reduced as the thrombosis blocked the vessel. The maximum velocity magnitude was 0.44 mm/s for the pre-stroke condition, and the speed decreased to nearly 0.1 mm/s in the during-stroke condition. Furthermore, the vessel was almost blocked in the post-stroke condition (Figs. 6(f)–(h)). In this demonstration, OSIV provided sufficient temporal resolution to observe the blood flow changes during the PT stroke injury in a mouse. We also show the dynamic OSIV maps for the three conditions in supplementary movie S2. Three representative OSIV velocity maps at 0.25 s intervals during the stroke are shown in supplementary Fig. S2.

**Fig. 6.**
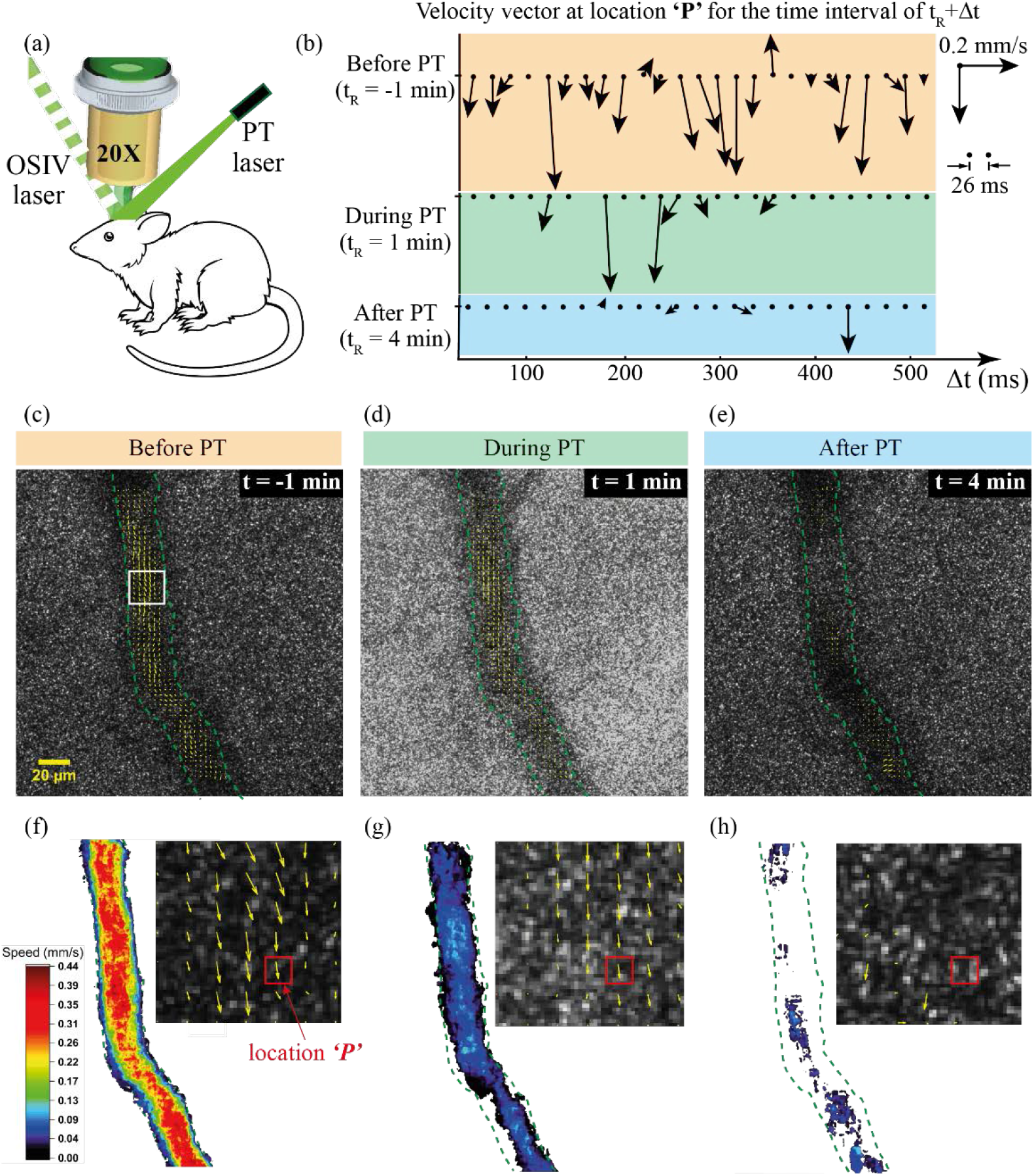
Application of OSIV in a photothrombosis stroke mouse model. (a) Schematic of the experiment. Two separate lasers were used for OSIV and photothrombosis global stroke. (b) OSIV measured velocity vector as a function of time at a location ‘P’ for the three condition of pre-stroke, during-stroke, and post-stroke. (c)–(e) The OSIV measured velocity map before, during, and after photothrombosis, respectively. (f)–(h) The OSIV measured flow speed map before, during, and after photothrombosis, respectively. Inset of each panel shows the magnified flow vector map of the boxed region in (c). The blood flow speed was the highest in the pre-stroke, then decreased during the photothrombosis and finally the vessel was blocked in the post-stroke condition.

## 4. Discussion

We developed a novel quantitative imaging technique—OSIV—to provide RBC velocity fields in a non-invasive, label-free, and wide-field manner by combining laser speckle physics and PIV. The current OSIV setup can measure RBC flow speed up to 7 mm/s, as determined from the solid diffuser experiment (Fig. 3). Therefore, the proposed technique is fast enough to detect the flow in cortical microvessels of a mouse such as capillaries and venules, where the mean flow speed is approximately 1.0 to 2.0 mm/s [37], and it can also detect the flow of blood in some arterioles [38]. Furthermore, we explored the feasibility of OSIV as a preclinical disease monitoring tool with a PT mouse model.

The main advantage of OSIV is the high velocity map refresh rate—48 map/s for the moving diffuser and the *in vitro* samples, which is greater than that of the video rate of 30 fps. For the *in vivo* case, the map refresh rate could be the same as that for the solid diffuser and fluidic phantom, that is, 48 map/s as shown in supplementary Movie. S2. This allows the study of wide-field functional brain activation, such as the hemodynamic responses with respect to various stimuli in the neurovascular coupling phenomenon [2]. OSIV is fast enough to resolve blood flow response that occurs within a few seconds. Hence, we applied OSIV to an animal ischemic stroke model to prove this concept. OSIV demonstrated real-time dynamic changes in the cortical blood flow field during during the occurrence of a stroke.

To implement a high OSIV map refreshing rate for real-time *in vivo* study, we utilized AOM on the illumination side and a high framerate sCMOS camera on the detection side. Modulated with a function generator, an AOM was used to optically chop the laser light for the illumination duration of 1.05 ms to reduce the speckle smearing with the shortest camera exposure time of 5.26 ms.

In this study, the frame rate was sufficient to prevent the integration of multiple decorrelated speckles into a single frame, as confirmed by the relatively high speckle contrast κ. To measure a higher blood flow by OSIV, one may use a faster camera commercially available, for example, the one used by Voleti et al. [39]. With the ROI of this camera (i.e., 1920 × 1080 pixels, which is larger than that of our sCMOS), the available maximum frame rate is 2,000 fps. In theory, OSIV can measure flow speed greater than 20 mm/s, which can allow the measurement of some arterial blood flow. In practice, to ensure a proper signal-to-noise ratio, laser power and camera frame rate need to be optimized within the tissue safety limit [40]. The limited field-of-view (FOV) demonstrated in the current study is due to the relatively slow camera (Andor Neo 5.5), and is not a fundamental limitation. We can increase the FOV by at least eight times with a faster camera [39], and thus OSIV can play an important role in a more clinically relevant setting.

Laser speckle contrast imaging (LSCI) [20] and multi-exposure speckle contrast imaging (MESI) [21] are contrast-based relative blood flow estimation methods that strictly rely on the ergodicity assumption. A comparison of our OSIV method with a common LSCI method is shown in supplementary Fig. S5. OSIV provides both quantitative blood flow speed and flow direction information, while LSCI shows only the contrast of the vessel with the remaining tissue (i.e., only qualitative flow speed and no flow direction information). Furthermore, LSCI did not give any physical unit for the relative blood flow index. To date, direct blood flow estimation has not been reported without assuming a scaling factor or fitting with theoretical models [20,21,23,41,42]. Therefore, a flow vector map cannot be obtained by these models. In comparison, OSIV can provide a direct RBC flow vector map without any calibration with flow phantom or using any complex model, as these methods are inaccurate when used in translational studies.

Critically, the development of OSIV paves a way to address the unmet clinical needs for continuous fast blood flow measurements with relatively simple instrumentation. However, there are still limitations of the method to validate the OSIV technique. We had to perform speckle imaging and white light imaging sequentially for the *in vivo* demonstration (supplementary Fig. S1). However, a coaxial illumination and detection path would allow simultaneous detection of white light imaging and speckle imaging.

For future development, a compact, portable, head-mounted OSIV instrument can allow functional brain imaging in awake mice [43]. Moreover, for faster OSIV, it would be interesting to incorporate deep learning [44], which may improve the accuracy of the velocity vector direction and reduce the number of averaging of correlation maps.

## 5. Acknowledgment, and disclosures

## 5.1. Acknowledgments

The work was supported by The GIST Research Institute (GRI) GIST-CNUH research collaboration grant funded by GIST in 2021 and the Joint Research Project of Institutes of Science and Technology in 2021. The National Research Foundation of Korea (N.R.F.) funded by the Korean government (MEST) (NRF-2019R1A2C2086003). The Brain Research Program through the N.R.F. funded by the Ministry of Science, I.C.T. & Future Planning (NRF-2017M3C7A1044964). The Ministry of Science and ICT, the Ministry of Trade, Industry and Energy, the Ministry of Health & Welfare, Republic of Korea, the Ministry of Food and Drug Safety (Project Number: 202011D13).

