## Supplementary Material for "Quantitative blood flow estimation *in vivo* by optical speckle image velocimetry"

### 1. Supplementary Methods

#### 1.1. Image Processing

The OSIV algorithms were implemented in MATLAB R2018. A two-dimensional median filter of 3×3 pixels was used in the post-processing to generate the OSIV maps. Each OSIV velocity map was obtained from the mean of four correlation maps, which results in an effective temporal resolution of 26.3 ms. The velocity map update rate was 48 map/s, which was higher than that of the typical video rate of 30 fps. To estimate the RBC speed between the two consecutive images from bright-field imaging, TrackMate [1] was used.

#### 1.2. Moving diffuser experiment

A solid glass diffuser (Thorlabs DG10-220) was placed on a sample holder (Thorlabs MP100-RCH1). The sample holder was placed on a motorized linear translation stage (SC-004-YF-0374, DCT Company Ltd., Korea), as shown in Fig. 2(b). The minimum speed was 2 mm/s. In the experiment, the diffuser was moved at speeds of 2 to 8 mm/s with an interval of 1 mm/s. The speckle images had a field-of-view of 332 μm × 332 μm (512 × 512 pixels).

#### 1.3. In vitro experiments

Five rats (Sprague Dawley, 10–14 weeks old with a body weight between 230 and 270 g) were used for blood collection. The animals were handled in accordance to the guidelines and instructions provided by the Institutional Animal Care and Use Committee (IACUC) at the Gwangju Institute of Science and Technology, South Korea. The experimental protocol was approved by the LARC GIST (Protocol Number: GIST-2020-083). We ensured that the time interval between the collection of blood from the same rat should be greater than two weeks. We collected a blood sample of 1 mL from the rat tail, and the collected sample was transferred to a blood collection tube (K2 EDTA (K2E), BD Vacutainer) [2]. The sample was washed with 1×PBS, and we prepared a 50% hematocrit solution using a centrifuge machine (Velocity 18R, Dynamica Scientific Ltd., UK) as detailed in a previous study [2].

The microchannel used in the fluid phantom was custom-manufactured by soft-lithography technology. The dimensions (length, width, and height) of the microfluidic channel were 20 mm, 1 mm, and 45 μm, respectively, as shown in supplementary Fig. S3. The diameters of the inlet and the outlet of the channel were 1 mm and 2 mm, respectively. Prior to injecting the blood sample into the microchannel, we pre-filled the channel with heparin (25,000 I.U./5mL) to avoid clotting. After a few minutes, the RBC sample was pushed into the microfluidic channel by a syringe pump (Pump11 Elite Harvard Apparatus, US). The conversion of flow rate into RBC speed is presented in the supplemental material table S4. The speckle image had a field-of-view of 332 μm × 332 μm (512 × 512 pixels). The ROI was selected at the center of the microfluidic channel.

#### 1.4. In vivo imaging of the mouse brain

The experiment was conducted using mice (C57BL/6, 8–12 weeks old with 18–23 g bodyweight). The animal handling was performed following the recommendation given by the guidelines of the IACUC of the Gwangju Institute of Science and Technology, South Korea. The Laboratory Animal Resource Center approved the protocol, Gwangju Institute of Science and Technology, South Korea (Protocol Number: GIST-2020-084). The mice were anesthetized with a Zoletil/Xylazine mixture in saline solution (60/10 mg/kg body weight), and the body temperature was maintained at 37.0–37.5 °C. With the cranial drilling diameter of 5 mm, a circular coverslip of 6 mm diameter was used to make the cranial window. We followed the craniotomy and imaging protocol from the procedure provided in the literature [2,3].

#### 1.5. Application of OSIV on the PT stroke model

In the PT stroke experiment, speckle images were acquired at pre-stroke, during-stroke, and post-stroke conditions. For the PT stroke generation, photosensitizer dye [4], Rose bengal dye (30000-5G, Sigma Aldrich) solution with a 20 mg/kg concentration was administered by bolus injection. For PT illumination, we used a 532 nm laser (MG-III-532 LD, Changchun New Industries Optoelectronics Technology Co., Ltd., China) with 20 mW on the window. The laser light was globally illuminated over the cranial window for 2 min, and at the start of the second minute, speckle images were captured. Speckle images for the pre-stroke condition were captured before the PT stroke. After 1 min of PT stroke, speckle images were captured for the during-stroke condition. After 2 min of PT stroke, speckle images were captured for the post-stroke condition.

#### 1.6. Comparison of OSIV with traditional speckle imaging

Common ways of imaging blood flow with laser speckle imaging techniques are laser speckle contrast imaging (LSCI) [5] and multi-exposure speckle imaging (MESI) [6]. We performed the comparison of our OSIV method with a widely used LSCI method as shown in Fig. S5. The OSIV method provides both quantitative blood flow speed and flow direction information, while LSCI shows only the contrast of the vessel with the remaining tissue (i.e. only qualitative flow speed and no flow direction information). Due to this reason, the relative blood flow index (rBFI) was commonly calculated by the following parameter in the LSCI community,

$$rBFI = \frac{1}{K^2} = \frac{\sigma^2}{\langle I \rangle^2}$$

where $K$ is the speckle contrast, $\sigma$ is the standard deviation, and $\langle I \rangle$ is the average intensity of the image. There is no unit for rBFI, which means it has an arbitrary unit; however, OSIV provides quantitative speed with the unit of mm/s, along with the flow direction information not possible with LSCI.

#### 3. Supplementary figures

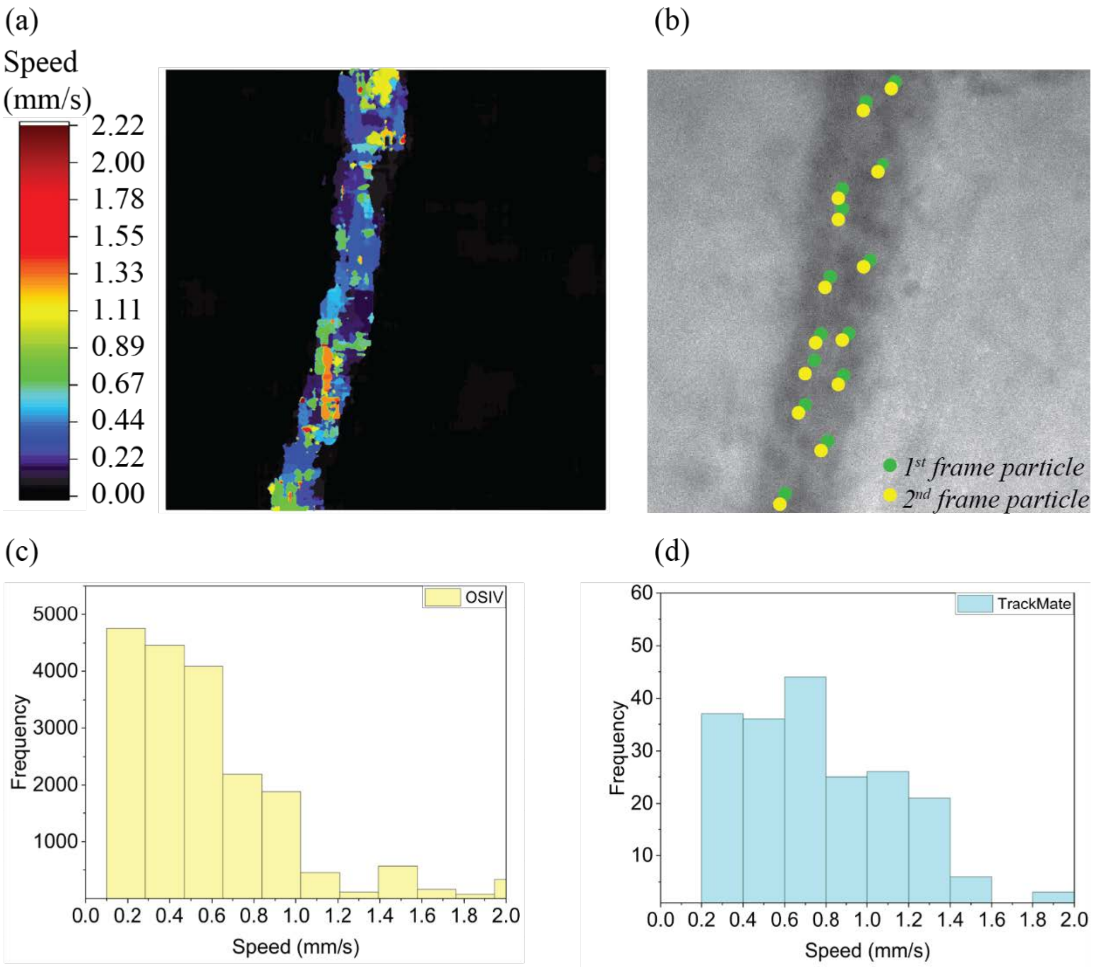


**Fig. S1. Comparison of OSIV speed map and white light image tagged with TrackMate.** (a) OSIV speed map for RBCs with the maximum value of 2.2 mm/s. (b) Two consecutive images were used in imageJ plugin 'TrackMate' for tracking the RBCs. The histogram of OSIV (c) have the similar range like the histogram (d) from the TrackMate.

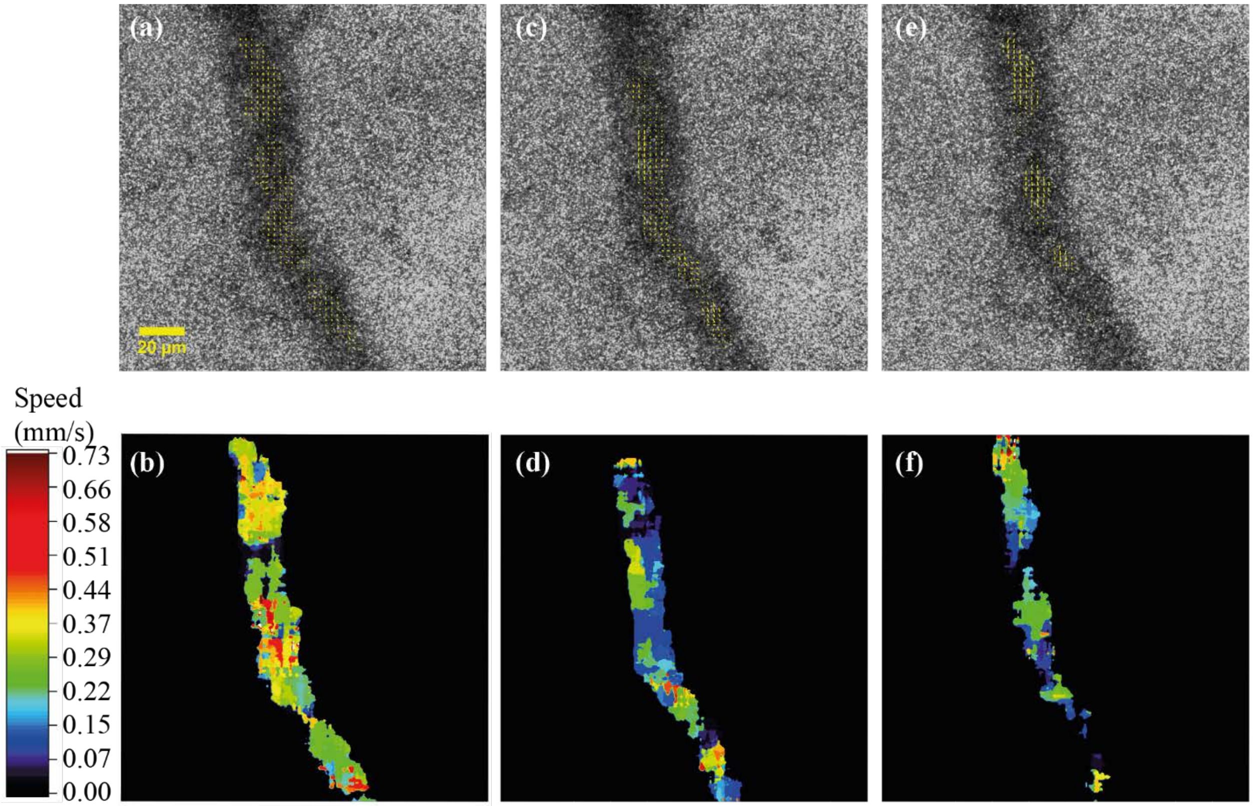


**Fig. S2. OSIV velocity field and magnitude maps during the stroke.** Three OSIV maps with a time interval of 0.25 s show that the overall RBC speed reduced.

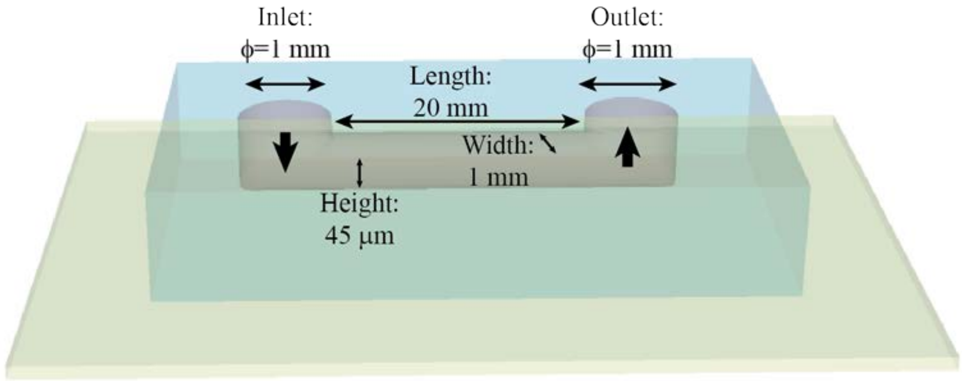


**Fig. S3. Schematic of the microfluidic channel.** The length, width and height of the microfluidic channel are 20 mm × 1 mm × 45 μm respectively. The inlet and the outlet of microfluidic channel are of 1 mm diameter.

| | Flow rate $'f'$ (µl/min) | Particle speed $v = {}^{f}/_{a}$ (mm/s) |
|---|---|---|
| 1 | 2 | 0.74 |
| 2 | 4 | 1.5 |
| 3 | 6 | 2.2 |
| 4 | 8 | 2.96 |
| 5 | 10 | 3.7 |

**Table S4. Conversion of flowrate in syringe pump to particle speed**. The first column shows the flowrate $'f'$ set on the syringe pump. The second column shows the particle speed $v$ in the microfluidic channel, and $'a'$ ($a = 0.045\ mm^2$) shows the cross-sectional area of the microfluidic channel.

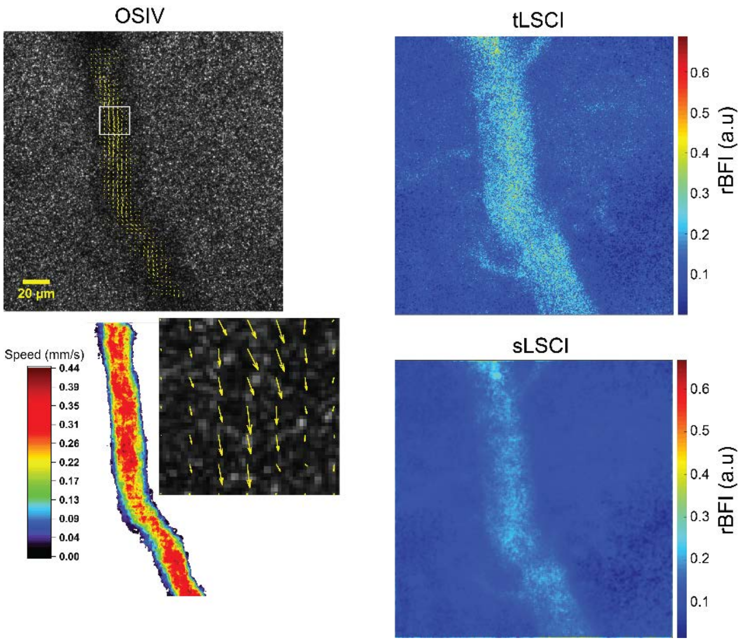


**Fig. S5. Comparison of OSIV and LSCIs for a blood vessel in a mouse brain.** Left: OSIV provides velocity (amplitude and direction) and speed map with unit of mm/s. Right: temporal LSCI (tLSCI) and spatial LSCI (sLSCI) provide the relative blood flow speed map with an arbitrary unit.
